# Interaction of Bunyamwera Virus Non-Structural Protein NSm with Cellular BNIP1 is Required for Efficient Viral Gene Expression and Replication

**DOI:** 10.64898/2026.07.01.735799

**Authors:** Rosalind F. Wartnaby, Juan Fontana, John N. Barr

## Abstract

Bunyamwera virus (BUNV) is the prototypical member of the *Peribunyaviridae* family of arthropod-borne viruses and possesses a genome comprising three segments of negative-sense RNA, named small, medium and large. The medium segment encodes a polyprotein that is processed to form Gn and Gc spikes and a non-structural protein, NSm. The role of NSm during replication in mammalian cells is poorly characterized, although it associates with a Golgi-derived structure called the virus factory (VF), the site of BUNV genome replication and virion assembly. To further define NSm function, we generated an epitope-tagged BUNV and used co-immunoprecipitation and quantitative proteomics to identify host interacting partners. NSm interacted with BCL-2 interacting protein 1 (BNIP1), a SNARE protein involved in COPI vesicle trafficking, with the importance of this interaction demonstrated by siRNA-mediated knockdown of BNIP1 expression, which significantly reduced BUNV gene expression and virion production. Interestingly, NSm also interacted with components of the NRZ complex, involved in COPI vesicle tethering in association with BNIP1, and inhibition of COPI complex formation resulted in loss of NSm expression. Taken together, our results identify BNIP1 as a host cell factor necessary for efficient BUNV replication and suggest the cellular localization of NSm at the VF is COPI-dependent.

## Introduction

The *Bunyaviricetes* class comprise segmented, negative-strand RNA viruses primarily transmitted by arthropod vectors (1, 2), and are capable of causing serious disease in humans, animals, and plants (1, 3). These include Crimean-Congo haemorrhagic fever virus, Lassa virus, and Rift Valley fever virus, all of which are listed by the World Health Organization as priority pathogens for research and development (4). In addition, the segmented nature of the bunyavirus genome enables reassortment, where segments are exchanged between co-infecting species, producing genetically distinct progeny and driving bunyavirus evolution through antigenic shift (5, 6). This process is a significant concern for public health, particularly when less harmful pathogens reassort to generate progeny with enhanced pathogenicity, exemplified by reassortment of non-pathogenic Bunyamwera (BUNV) and Batai orthobunyaviruses to generate Ngari virus (NRIV), which is associated with severe haemorrhagic fever in humans (7). Despite the threat posed by bunyaviruses, there are currently no clinically approved vaccines or antiviral therapeutics available for the prevention and treatment of human infections (8).

BUNV is classified within the *Orthobunyavirus* genus of the *Peribunyaviridae* family and provides a useful research model as a safer alternative to more pathogenic orthobunyaviruses such as NRIV, with lower biosafety level requirements for its study (1, 9). The BUNV genome consists of three RNA segments termed small (S), medium (M), and large (L), corresponding to their relative sizes (10). On overlapping reading frames, the S segment encodes NP and the non-structural protein NSs, an antagonist of the host cell interferon response (11, 12). The M segment encodes a glycoprotein precursor (GPC) which is co-translationally cleaved to form two glycoproteins, Gn and Gc, and the non-structural protein NSm. Gn and Gc form glycoprotein spikes inserted into the viral envelope that are required for virus entry (13), while the function of NSm is poorly characterized, although a role in penetration of the mid-gut has been recently described during the insect phase of the BUNV lifecycle (14). Finally, the L segment encodes the vRdRp, necessary for viral genome replication and transcription (15). All segments are protected by encapsidation by the nucleocapsid protein (NP) (16), and together with the viral RNA-dependent RNA polymerase (vRdRp), form viral ribonucleoprotein complexes (vRNPs) that are surrounded by host cell Golgi-derived lipid envelopes to form infectious virions (17).

NSm is a 175 amino acid-long integral membrane protein, which is cleaved from the GPC by ER-resident signal peptidase (SP, (18)). NSm has five identified domains, of which domains I, III and V are predicted to be transmembrane, with domain II likely extending into the Golgi lumen, and domain IV being cytosolic (19). NSm is cleaved from the GPC by virtue of SP recognition sequences in domains I and V (18) and subsequently localizes to the Golgi where it is reported to associate with a structure known as the virus factory (VF, (17)).

The VF is a structure created by the virus within the host cell where genome replication or virion assembly occur, often involving reorganization of cellular endomembranes, recruitment of host cell organelles and accumulation of viral components that together facilitate efficient virus production (20). VFs form across a wide range of intracellular sites for different viruses (21). In mammalian cells, the BUNV VF forms around the Golgi complex, inducing the formation of characteristic structures known as viral tubes, as well as recruiting mitochondria and membranes derived from the rough endoplasmic reticulum (RER, (17)).

Despite its localization and apparent involvement in VF formation within mammalian cells, the role of the BUNV NSm protein is unclear. The NSm protein is able to tolerate sizable deletions and insertions, except for the N-terminal domain, which is reported to be required for BUNV morphogenesis. Thus, it has been suggested that the BUNV NSm is mainly dispensable for mammalian infections (14), with similar findings for NSm proteins of other bunyaviruses (22–25). However, studies with NSm deletion mutants targeting alternative domains of NSm have indicated that while non-essential, this non-structural protein is necessary for efficient BUNV replication, and an important factor in VF functions (17, 18, 19). While a role for NSm has been recently uncovered in cell-to-cell spread and egress from the mosquito midgut barrier during the arthropod phase of the BUNV lifecycle (14), the precise function of NSm at the Golgi-associated VF in mammalian cells remains unknown.

Here, we investigated the role of NSm during mammalian BUNV infection. We generated a recombinant BUNV (rBUNV) variant harboring a hemagglutinin (HA) epitope tag in NSm, and used this tool to localize NSm to the VF in infected cells as well as perform co-immunoprecipitation and quantitative proteomics analysis to identify NSm-host cell interaction partners. One such interactor was soluble N-ethylmaleimide-sensitive factor attachment protein receptor (SNARE) protein BCL-2 interacting protein 1 (BNIP1), with this interaction validated using immunological methods. Visualization by immunofluorescence in infected cells revealed BNIP1 shifted from a diffuse cytoplasmic distribution in uninfected cells to a concentrated perinuclear region upon infection, colocalizing with NSm, while an siRNA knockdown of expression revealed BNIP1 to be necessary for efficient BUNV gene expression and virus production. NSm also interacted with components of the NRZ complex, involved in COPI vesicle tethering in association with BNIP1, and inhibition of COPI complex formation abolished NSm expression. Taken together, these findings identify BNIP1 as an NSm interaction partner and a host cell factor necessary for efficient BUNV replication and suggest that NSm localization to the VF is dependent on COPI-mediated trafficking.

## Results

### NSm is required for efficient BUNV replication in mammalian cells

To determine the necessity of NSm in mammalian infections, we generated rBUNVΔNSm (18), a recombinant BUNV lacking the full mature NSm (domains II-V), while retaining the N-terminal NSm signal peptide (NSm domain I) and cleavage sites necessary for GPC processing (Fig. 1A, (18, 19)). Firstly, the plaque phenotypes of rBUNV-WT (wildtype) and rBUNVΔNSm were compared (Fig. 1B). In agreement with previous reports (18), rBUNVΔNSm exhibited a smaller plaque phenotype than rBUNV-WT, indicating that deletion of NSm resulted in attenuation of virus growth. Next, the peak virus yields of the two viruses were compared (Fig. 1C and supplementary fig. S1). A549 cells were infected with either rBUNV-WT or rBUNVΔNSm at a multiplicity of infection (MOI) of 0.1, before viral supernatants were harvested at 48 hours post-infection (hpi), the timepoint of maximum BUNV titer, for subsequent determination of viral titers by plaque assays. To enable accurate quantification, rBUNVΔNSm plaque assays were incubated for an additional 24 hours compared to rBUNV-WT to allow visible plaque formation. Consistent with prior findings (18, 19), replication of rBUNVΔNSm was compromised compared to rBUNV-WT, evidenced by a significant reduction of maximum virus yield (Fig. 1C). These results indicate that NSm is required for optimal BUNV replication in mammalian cells.

**Fig 1.**
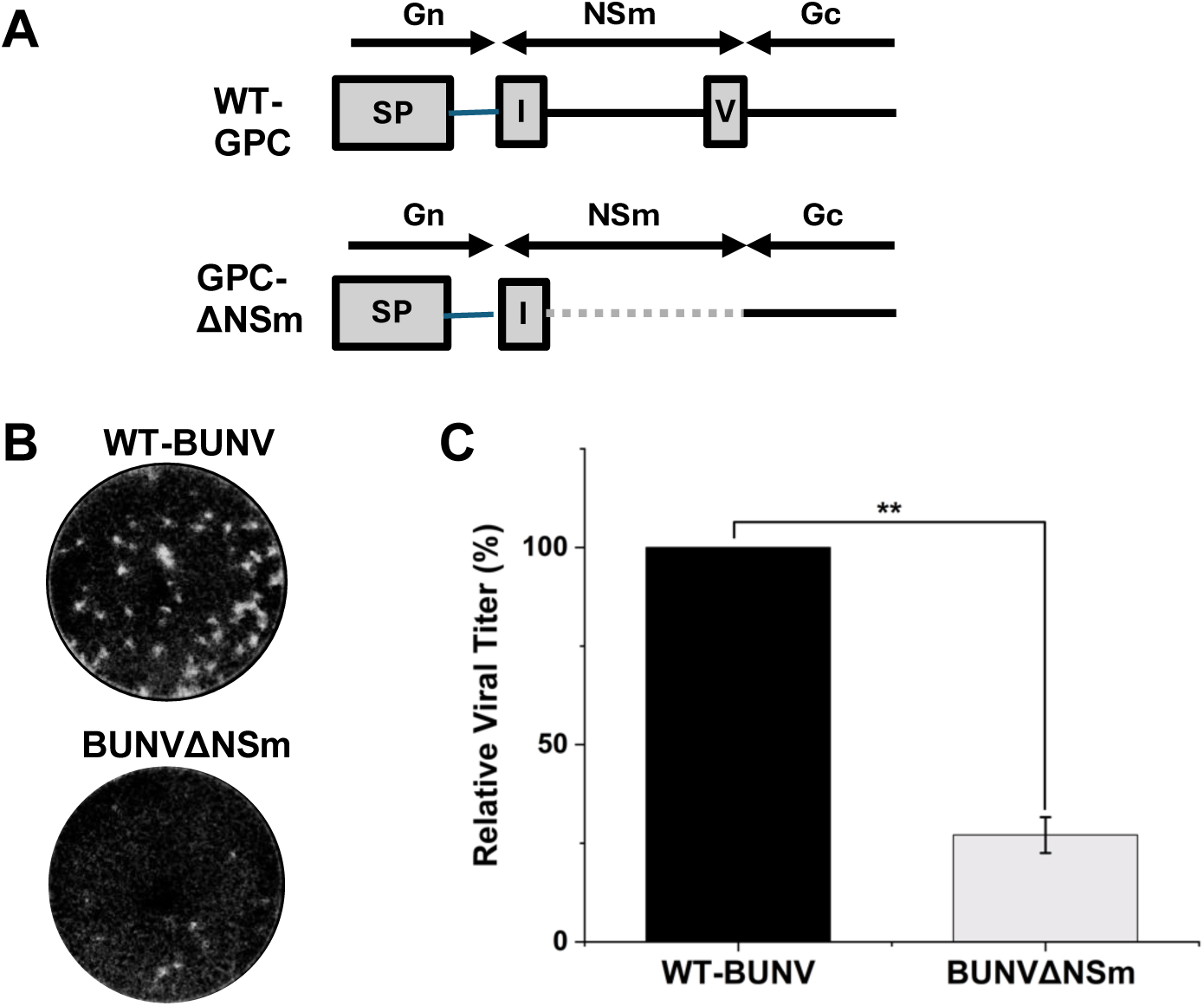
Comparison of the growth kinetics of rWT-BUNV and rBUNVΔNSm. (**A**) Schematic showing the deletion of NSm domains II-V from GPC in rBUNVΔNSm compared to WT-GPC, with signal peptides (SPs) represented by grey boxes. (**B**) Comparison of the plaque phenotypes of rWT-BUNV and rBUNVΔNSm at three days post-infection (dpi), demonstrating a smaller plaque phenotype for rBUNVΔNSm. (**C**) Comparison of mean peak virus yields indicating attenuation of rBUNVΔNSm – A549s were infected with rWT-BUNV or rBUNVΔNSm at an MOI of 0.1, before viral supernatants were harvested at 48 hpi for subsequent determination of viral titers by plaque assays, with normalization to rBUNV-WT and three biological replicates performed. Plaque assays were incubated for four days in the case of rBUNVΔNSm compared to three days for rBUNV-WT to allow visible plaque formation for quantification. Error bars indicate mean ± standard deviation. Representative plaque assay images are displayed in Fig. S1.

### Generation of a recombinant BUNV variant harbouring an HA tag in NSm

As deletion of NSm compromises efficient BUNV replication, and NSm has been implicated in BUNV VF function (17), we wished to generate a tool to visualize NSm in infected cells and uncover interactions between NSm and host cell proteins that might elucidate the role of NSm at the VF. To achieve this, we produced a recombinant BUNV harboring an HA tag inserted into NSm (rBUNV-NSm-HA). To choose the tag insertion site, AlphaFold 3 (26) was used to predict the structure of the mature NSm protein (Fig. 2A), revealing two unstructured regions, consistent with previously identified luminal and cytoplasmic domains II and IV (Fig. 2B, (18, 19)). Due to its ability to tolerate deletions (19), and to ensure the best accessibility for antibodies, cytoplasmic domain IV was selected. Subsequently, rBUNV-NSm-HA was successfully rescued, with resulting plaque assays (Fig. 2C) and western blotting (Fig. 2D and supplementary fig. S2) confirming the generation of infectious virus with a titer of approximately 10^7^ PFU/mL.

**Fig 2.**
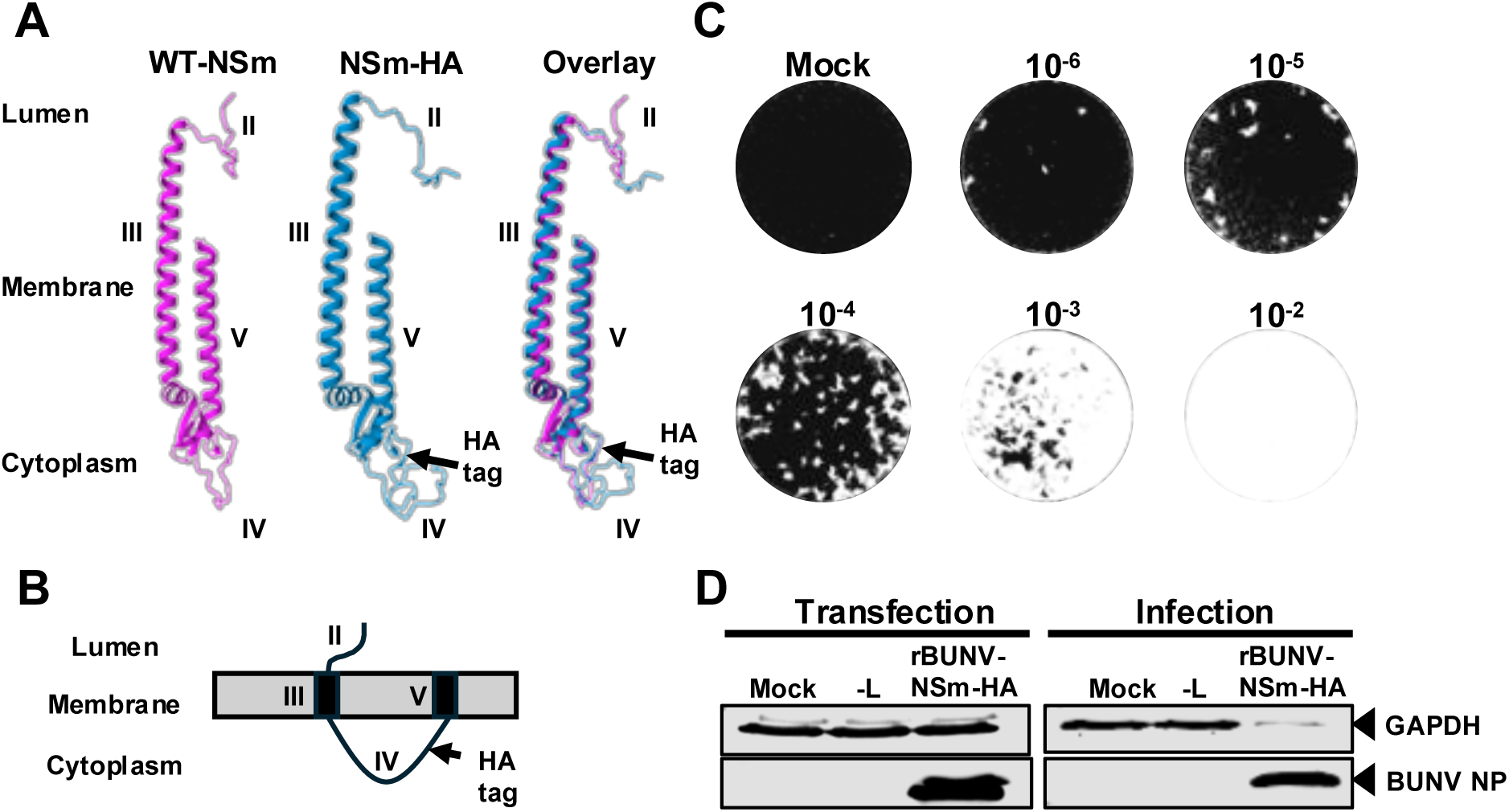
Generation of a recombinant BUNV variant harboring an HA epitope tag in NSm. (**A**) AlphaFold 3 (26) structural predictions of mature WT-NSm and NSm-HA, with the HA tag inserted into the cytoplasmic domain IV of NSm. (**B**) The predicted topology of the mature NSm protein, with the HA tag location indicated (18, 19). Hydrophobic domains are indicated by black columns spanning the intracellular membrane, indicated by the grey rectangle. (**C**) Plaque assay for rBUNV-NSm-HA demonstrating the presence of plaques at three dpi – ten-fold serial dilutions of supernatant from infected cells were created with dilution factors indicated and used to infect BHK-21 cells. (**D**) Western blot analyses of BSR-T7 cell lysates transfected with rescue plasmids alongside (-) L and mock transfected cells, and BHK-21 cell lysates infected with the transfection supernatants, with GADPH used as a loading control, with the presence of BUNV NP used to confirm successful virus rescue. Uncropped western blots are displayed in Fig. S2.

### Localization of NSm and NP in infected cells

To assess the localization of NSm and NP, and to evaluate the accessibility of the tag for successful antibody detection, we infected A549 cells with rBUNV-WT or rBUNV-NSm-HA at an MOI of 0.1 before performing immunofluorescence staining at 24 hpi to detect NP and HA (Fig. 3A). While NP staining was distributed throughout the cytoplasm, the NSm signal was condensed in the perinuclear region to one side of the nucleus, likely indicating the presence of NSm at the Golgi or ER. To provide additional information regarding the subcellular location of NSm in infected cells, we next visualized NSm alongside cellular markers previously identified as components within the BUNV VF (17). A549 cells were infected with rBUNV-NSm-HA at an MOI of 0.5 and stained at 24 hpi for the *cis*-Golgi marker Golgi Matrix protein 130 kDa (GM130), mitochondrial marker Translocase of Outer Mitochondrial Membrane 20 (TOM20), and rough ER marker calnexin (CANX), alongside the HA tag of NSm (Fig. 3B). These cell markers were found to localize to the same perinuclear area as NSm, with particularly notable colocalization observed between NSm and GM130, a *cis*-Golgi marker, through line scan analysis (Fig. 3C), consistent with previous reports describing NSm at the Golgi-derived VF (17).

**Fig 3.**
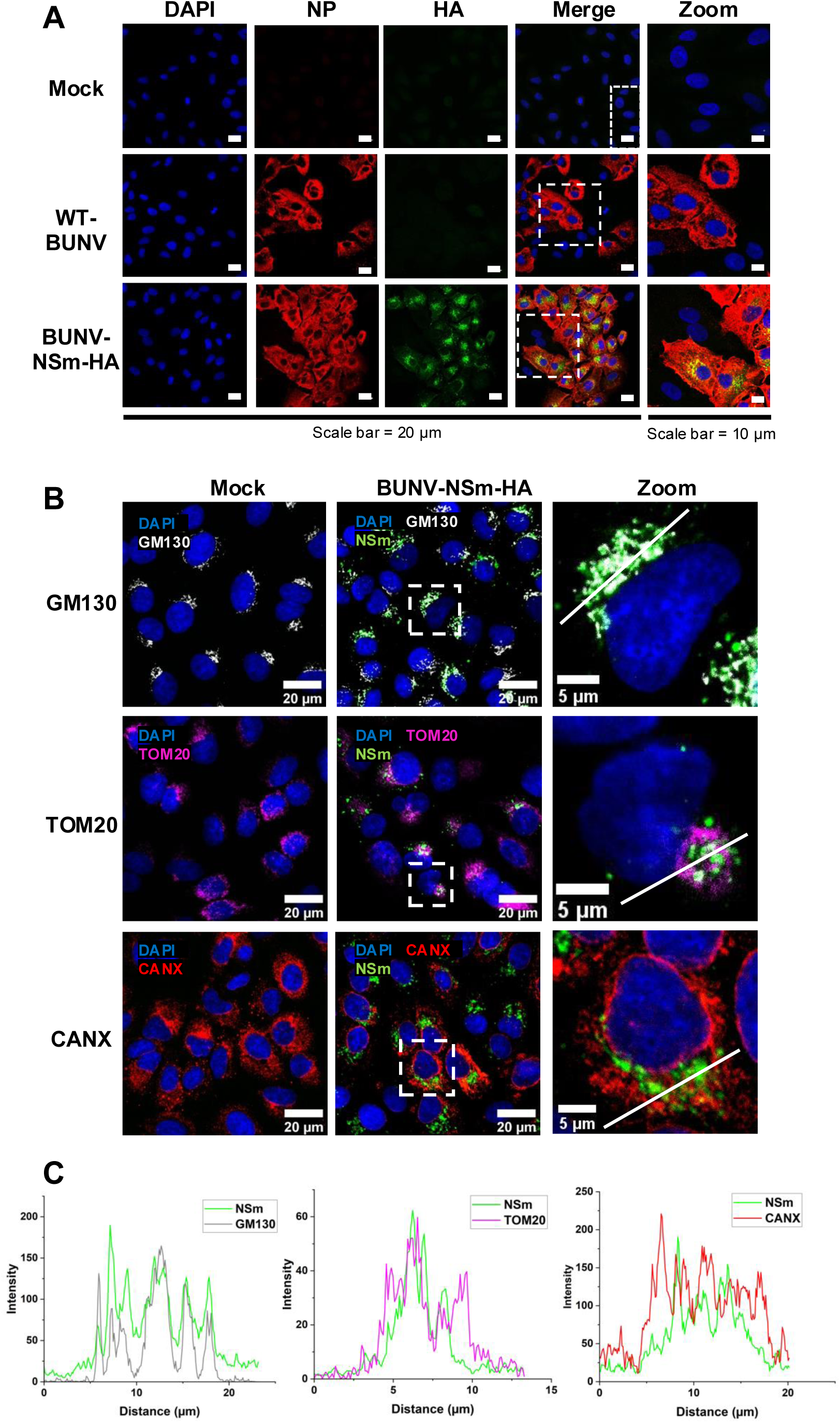
Localization of NSm in BUNV-infected cells. (**A**) Confocal microscopy images of uninfected, rWT-BUNV-infected, and rBUNV-NSm-HA-infected A549 cells at 24 hpi at an MOI of 0.1, with staining to detect NSm, NP, and DAPI, demonstrating good accessibility and detection of the HA tag. (**B**) Confocal microscopy images of uninfected and rBUNV-NSm-HA-infected A549 cells at 24 hpi at an MOI of 0.5, with co-staining for NSm and DAPI alongside GM130, TOM20, or CANX. (**C**) Line scan analyses for the regions indicated in panel (B), displaying the relative intensities of NSm with GM130, TOM20, and CANX, revealing strong colocalisation between NSm and GM130.

### Co-immunoprecipitation and quantitative proteomics to identify NSm-host cell interaction partners

In order to gain insights into the function of NSm and its potential involvement in the BUNV VF in mammalian cells, we wished to determine interactions between NSm and host cell proteins. To achieve this, we infected A549 cells with rBUNV-NSm-HA at an MOI of 0.5, alongside a non-infected condition, before harvesting cell lysates at 24 hpi. Next, we performed a co-immunoprecipitation, binding HA antibody to the surface of protein G magnetic beads and using the HA tag to isolate NSm and any interacting partners from the rBUNV-NSm-HA-infected cell lysate. Following this, interacting proteins were identified through liquid chromatography-tandem mass spectrometry (LC-MS/MS, supplementary data set S1). The resulting proteins were filtered through the removal of any hits present in the non-infected mock sample that bound non-specifically with any measurable peak area, as well as those identified as likely laboratory contaminants such as human keratins, yielding 420 host cell proteins associated with NSm. The remaining proteins were ranked by peak area values, providing an estimation of relative protein abundance based on the chromatographic signal intensity, alongside consideration of peptide-spectrum match (PSM) values, which reflect how many MS/MS spectra were assigned to peptides from each protein, and unique peptide counts, indicating how many peptide sequences map uniquely to a given protein. Together, these metrics were used to prioritize high-confidence and abundant interaction partner candidates (Supplementary table S1).

### Independent validation of the NSm-BNIP1 interaction

One of the top host cell interaction partners identified through the mass spectrometry analysis was BNIP1, a SNARE protein involved in the transport of coatomer protein complex 1 (COPI)-coated vesicles (27). To substantiate this result, we wished to immunologically validate the interaction between NSm and BNIP1. Firstly, we used western blotting to probe the mock and infected input cell lysates and bead-bound co-immunoprecipitation products for NSm and BNIP1 (Fig. 4A and supplementary fig. S3). As expected, an NSm band was observed for the infected lysate and co-immunoprecipitation product, but not for the mock counterparts, indicating the successful capture of NSm on the beads. For BNIP1, a prominent band was detected in the bead-bound co-immunoprecipitation product from the infected lysate, but not for the mock equivalent, demonstrating a specific interaction between NSm and BNIP1.

**Fig 4.**
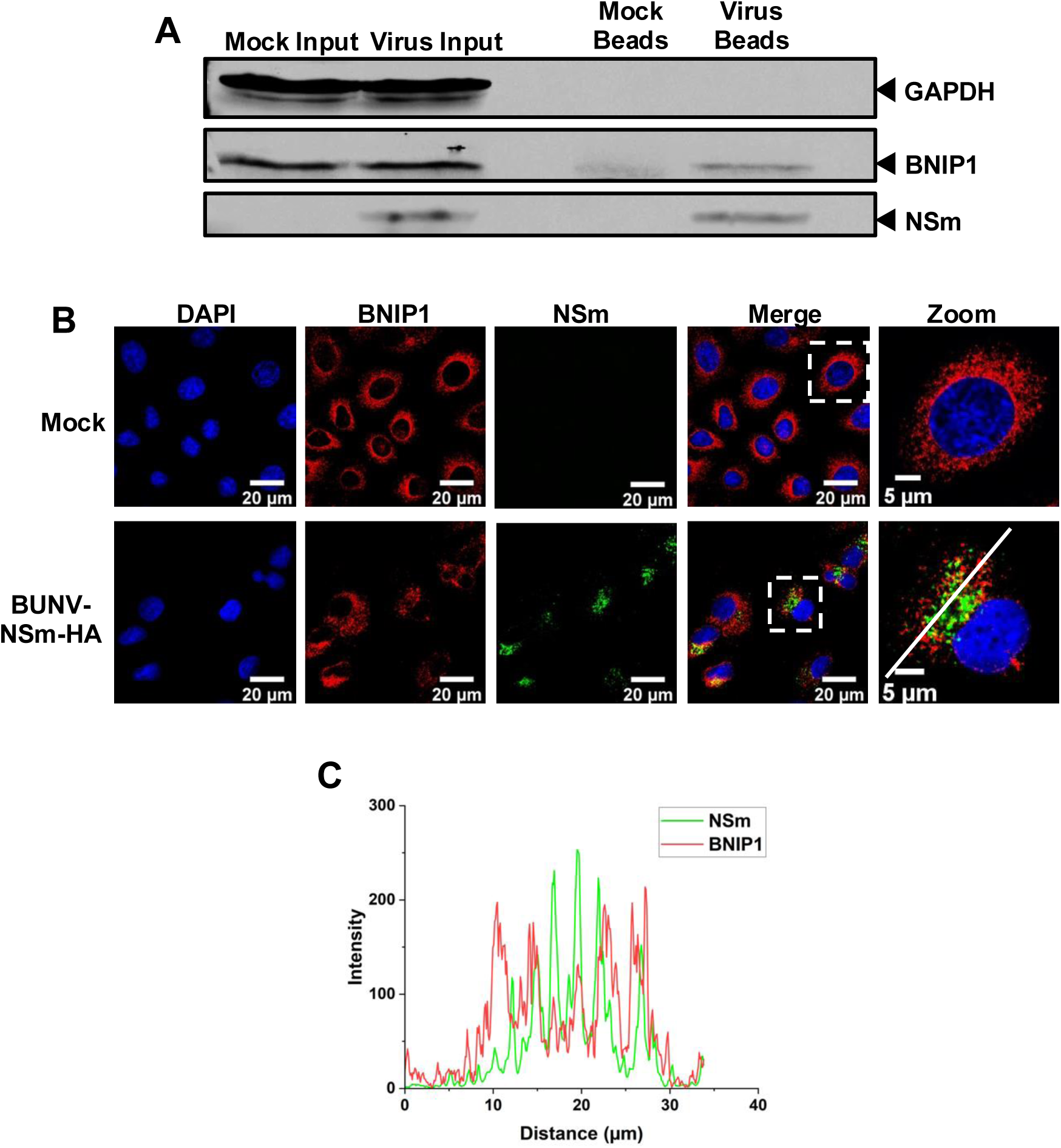
Analysis of the interaction between NSm and BNIP1 in mammalian cells. (**A**) Western blot analysis of NSm immunoprecipitated from rBUNV-NSm-HA-infected A549 cell lysate, with probing for NSm and BNIP1 alongside GAPDH as a loading control. This shows BNIP1 to be co-immunoprecipitated with NSm. The uncropped western blot is displayed in Fig. S3. (**B**) Confocal images of uninfected or rBUNV-NSm-HA-infected A549 cells at 24 hpi at an MOI of 0.5, with staining to detect NSm, BNIP1, and DAPI, demonstrating redistribution of BNIP1 in infected cells and its colocalisation with NSm. (**C**) Line scan analysis of the region indicated in panel (B), indicating the relative intensities of the NSm and BNIP1 channels along the scan line.

Secondly, we assessed colocalization of NSm and BNIP1 signals through immunofluorescence staining of cells infected with rBUNV-NSm-HA at an MOI of 0.5 at 24 hpi using HA and BNIP1 antisera (Fig. 4B). NSm and BNIP1 were found to localize to the same perinuclear region of the cell, with line scan analysis demonstrating alignment of several of the peak intensities of the two proteins (Fig. 4C), supportive of an interaction between NSm and cellular BNIP1. Additionally, a change in distribution of BNIP1 within the cell was observed upon BUNV infection, with a more diffuse dispersion of BNIP1 staining throughout the cytoplasm in uninfected cells, compared to a more compact signal at perinuclear sites in infected cells (Fig. 4B), suggesting a potential recruitment of BNIP1 to or around the VF during BUNV infection.

### Cellular BNIP1 is required for efficient viral gene expression and replication

To further evaluate the importance of BNIP1 during the BUNV replication cycle, we performed an siRNA-mediated knockdown of BNIP1 in A549 cells to assess how reduced expression of this protein affected BUNV gene expression and replication, processes that occur at the VF (Fig. 5A and B and supplementary fig. S4). Briefly, A549 cells were reverse transfected with siRNAs specific for BNIP1, alongside both scrambled siRNA and no siRNA negative controls, with knockdown allowed to progress for 24 hours. Following a short recovery period, cells were infected with rBUNV-WT at an MOI of 0.5, before viral supernatants and cell lysates were harvested at 24 hpi. To test for changes to BUNV gene expression, western blotting was used to probe for the BUNV NP protein (Fig. 5C and D and supplementary fig. S4). To test for changes to BUNV multiplication, viral titers were calculated through plaque assays (Fig. 5E and supplementary fig. S4). Compared to control cells subjected to transfection with either scrambled or no siRNAs, knockdown of BNIP1 resulted in significant decreases in both NP protein expression and viral titers, demonstrating that cellular BNIP1 is necessary for efficient BUNV gene expression and replication.

**Fig 5.**
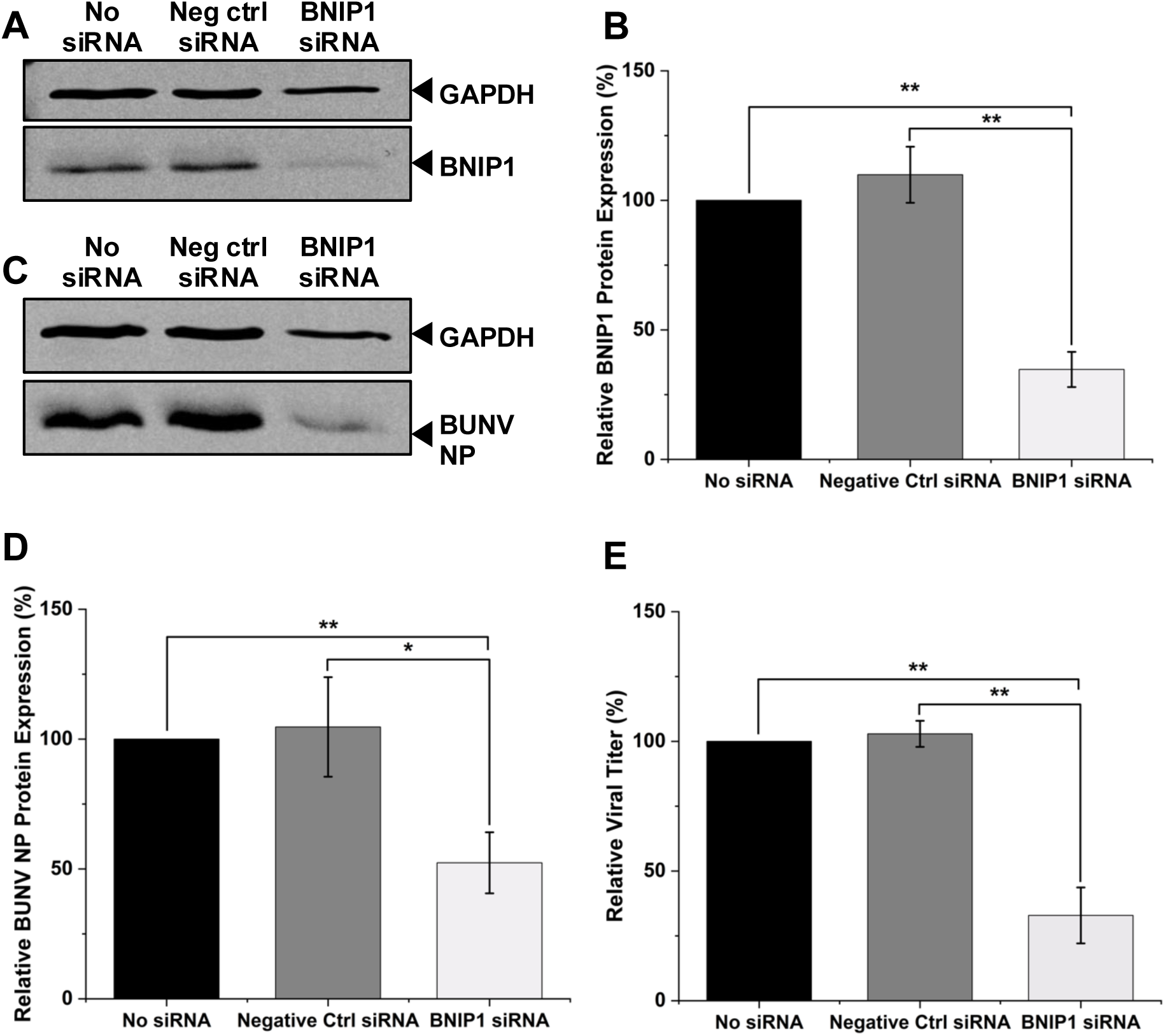
Effects of siRNA-mediated knockdown of host cell BNIP1 on BUNV gene expression and replication. A549 cells were reverse-transfected with siRNA, before infection with rBUNV-WT at an MOI of 0.5, and harvesting of cell lysates and viral supernatants at 24 hpi. Three biological replicates were performed with the mean results displayed on graphs, and results normalized to the no siRNA control. Error bars indicate mean ± standard deviation. Uncropped western blots for all replicates are displayed in Fig. S4A. (**A**) Western blot analysis of BNIP1 alongside GAPDH demonstrates a successful knockdown of BNIP1 protein levels in cells transfected with siRNA targeting BNIP1. (**B**) Densitometry analysis indicating successful knockdown of BNIP1 protein levels in cells transfected with siRNA targeting BNIP1 across all replicates. (**C**) Western blot analysis of BUNV NP alongside GAPDH reveals a reduction in NP protein levels when host cell BNIP1 is knocked down. (**D**) Densitometry analysis demonstrating a decrease in BUNV NP protein levels when BNIP1 is knocked down across all replicates. (**E**) Viral titers of supernatants across the replicates determined by plaque assays demonstrate a reduction in viral titer when BNIP1 is knocked down. Representative plaque assay images are displayed in Fig. S4B.

### NSm interacts with the NRZ complex, which associates with BNIP1 and facilitates COPI vesicle trafficking

ER t-SNAREs, including BNIP1, are known to associate with the mammalian NRZ tethering complex, composed of Neuroblastoma Amplified Sequence (NBAS), Zeste White 10 (ZW10), and RAD50 Interactor 1 (RINT1, (28, 29)). Together, these proteins are thought to facilitate the tethering and fusion of COPI-coated vesicles with their target membrane, as reported for the yeast homolog Dsl1 complex (30–32). Given the importance of BNIP1 in efficient BUNV infection, we next investigated whether any other components of this machinery also interact with NSm. Our mass spectrometry analysis revealed that in addition to BNIP1, two of the three NRZ complex components, NBAS and ZW10, were uniquely pulled down by NSm. To further validate the interaction between NSm and the NRZ complex, we performed western blotting to probe for the presence of ZW10 in the bead-bound pull-down products following the NSm co-immunoprecipitation, alongside mock and infected input cell lysates (Fig. 6A and supplementary fig. S5). ZW10 protein expression was observed in the infection co-immunoprecipitation product, but not in the uninfected counterpart, corroborating a specific interaction between NSm and ZW10.

**Fig 6.**
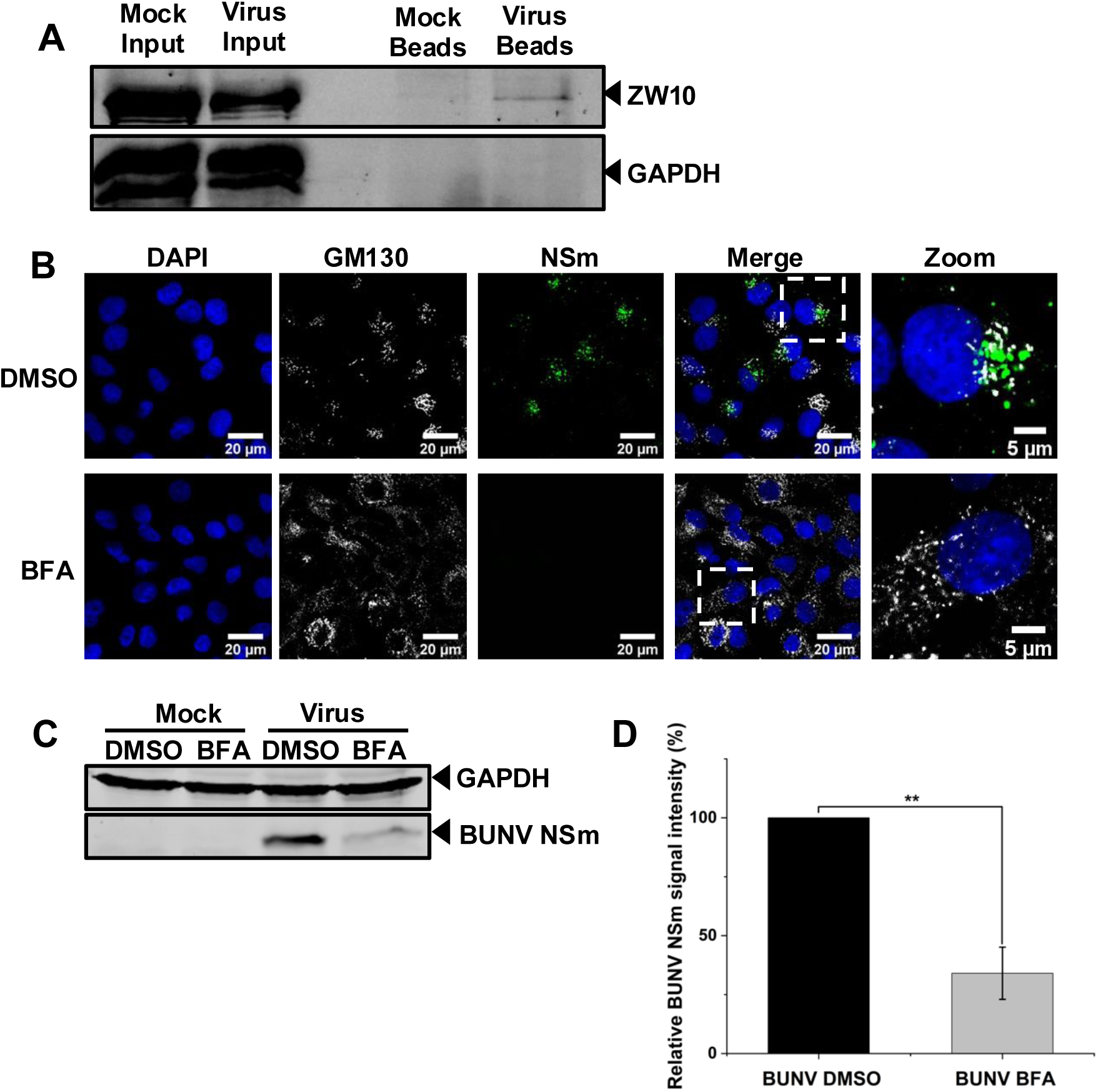
Analysis of the NSm-ZW10 interaction and the effect of COPI inhibition on NSm protein expression during BUNV infection in mammalian cells. (**A**) Western blot analysis of NSm immunoprecipitated from rBUNV-NSm-HA-infected A549 cell lysate, with probing for ZW10 alongside GAPDH as a loading control revealing ZW10 to be co-immunoprecipitated with NSm. The uncropped western blot is displayed in Fig. S5A. (**B-D**) A549 cells were infected with rBUNV-NSm-HA at an MOI of 5, before treatment with DMSO or 5 µg/mL BFA at 1 hpi. (**B**) Confocal images of cells at 8 hpi with staining to detect NSm, GM130, and DAPI, demonstrating a reduction of NSm signal in BFA-treated cells. (**C**) Western blot analysis of cell lysates collected at 8 hpi with probing for BUNV NSm alongside GAPDH, revealing a reduction in NSm protein levels in BFA-treated cells. The uncropped western blot is displayed in Fig. S5B. (**D**) Densitometry analysis demonstrating a decrease in BUNV NSm protein levels in BFA-treated cells across three biological replicates, with the mean results displayed, and normalized to the DMSO control. Error bars indicate mean ± standard deviation.

### BFA treatment reduces NSm abundance in cells infected with BUNV

Due to the interaction of NSm with components involved in COPI vesicle trafficking, we wished to further investigate the potential involvement of COPI during BUNV infection. To achieve this, we used Brefeldin A (BFA), which disrupts COPI-dependent trafficking by inhibiting ARF1 activation, thereby preventing recruitment of the COPI coat complex to Golgi membranes and causing its dissociation from the Golgi (33, 34). A549 cells were first infected with rBUNV-NSm-HA for one hour prior to treatment to avoid confounding effects on virus entry. Subsequently, cells were treated with 5 µg/mL BFA, a concentration previously determined as non-toxic in our human-origin A549 cells (35), before being fixed or lysed at 8 hpi. To assess any changes to BUNV NSm gene expression, immunofluorescence staining (Fig. 6B) and western blotting were used to probe for BUNV NSm protein expression (Fig. 6C and D and supplementary fig. S5). Compared to control DMSO-treated cells, treatment with BFA resulted in a reduction of NSm localization at the Golgi as well as a significant decrease in overall NSm protein expression, suggesting that the cellular localization of NSm at the Golgi-associated VF is COPI-dependent.

## Discussion

A role has recently been described for NSm during the arthropod phase of the BUNV lifecycle, where a requirement for NSm for successful cell-to-cell spread and release from the mosquito midgut was identified (14). Here, we use NSm deletion mutants to demonstrate that NSm is also important for efficient virus replication in mammalian cells, in agreement with previous findings (17, 18, 19).

NSm has been previously detected at the VF during mammalian infection (17). Formed at the Golgi complex, the BUNV VF plays a critical role in genome replication and virus assembly (17) and provides a likely location of reassortment during co-infections. The use of different membranous compartments within the host cell to facilitate assembly is characteristic of enveloped viruses, including other RNA viruses such as coronaviruses and flaviviruses (36–40). Despite its association with the VF and apparent involvement in VF formation within mammalian cells, the exact role of NSm is unknown.

In this study, we aimed to address this question through the identification of NSm-host cell protein interaction partners. Using a recombinant BUNV variant harboring an epitope-tagged NSm, we performed a co-immunoprecipitation and quantitative proteomics, revealing an interaction between NSm and SNARE protein BNIP1. Immunological analyses of this interaction, alongside an siRNA-mediated knockdown of BNIP1 expression in the host cell, corroborated BNIP1 as a contributing host cell factor required for efficient BUNV gene expression and virus production.

Firstly, we used rBUNVΔNSm (18) to demonstrate that NSm is required for efficient BUNV replication, concurring with previous studies using NSm deletion mutants (18, 19). To learn more about the involvement of NSm during mammalian BUNV infections, we designed and generated rBUNV-NSm-HA, a recombinant BUNV variant containing an HA epitope tag in NSm. This facilitated the visualization of NSm in infected cells, revealing its localization to the perinuclear area of the cell, in contrast to a more dispersed distribution of NP throughout the cytoplasm. Strong colocalization was observed between NSm and GM130, a *cis*-Golgi marker, supporting the presence of NSm at the Golgi-derived VF, and a potential role for NSm in VF functions. This result is in agreement with previous findings that NSm localizes to the VF, where it may play a role in maintaining the structure of viral tubes harbouring viral replication complexes (17).

Next, we aimed to learn more about the function of NSm during mammalian infection by using rBUNV-NSm-HA to perform a co-immunoprecipitation and quantitative proteomics to identify interactions between NSm and host cell proteins. BNIP1 was identified as a top host cell protein interactor, with independent validation of the interaction through western blot analysis of co-immunoprecipitation products. BNIP1 is an endoplasmic reticulum (ER)-resident target SNARE (t-SNARE), a component of a SNARE complex thought to be involved in the retrograde transport of COPI-coated vesicles from the Golgi to the ER (27). This is consistent with the established function of Sec20, the yeast homolog of BNIP1, in this retrograde transport route (41). SNARE proteins have also been implicated in the replication and egress of other RNA viruses, such as paramyxoviruses, tombusviruses, and enteroviruses (42–44). In addition to its function as a t-SNARE, BNIP1 has been suggested to play a role in regulating apoptosis (45–47), as well as influencing mitochondrial dynamics (48).

Assessment of the cellular distribution of BNIP1 and NSm during BUNV infection using immunofluorescence showed that while BNIP1 expression was more diffuse throughout the cytoplasm in uninfected cells, its signal was concentrated in the perinuclear region in infected cells and colocalized with NSm, suggesting a potential recruitment of BNIP1 to the Golgi-associated VF by NSm during infection. To further evaluate the importance of BNIP1 during BUNV infection, we performed an siRNA-mediated knockdown of host cell BNIP1, which revealed BUNV gene expression and virus production to be significantly compromised when BNIP1 levels were depleted. Taken together, these findings suggest that BNIP1 is a host cell factor that interacts with NSm and plays an important role in supporting efficient viral gene expression and replication.

ER t-SNAREs including BNIP1 interact with the NRZ complex, a CATCHR (complexes associated with tethering containing helical rods) complex composed of subunits NBAS, RINT1, and ZW10 (28, 29). Together, these interactions are thought to help mediate the tethering and fusion of COPI-coated vesicles with their target membrane, as seen in its yeast counterpart Dsl1 complex (30–32). The involvement of BNIP1 in this process is further substantiated by the recruitment of BNIP1 into COPI-coated vesicles (27). In their native cellular role, COPI vesicles are thought to mediate the retrograde transport of proteins and lipids from the Golgi to the ER, as well as the bi-directional movement of cargo between different Golgi compartments (49). Due to the importance of BNIP1 during BUNV infection, we wished to determine whether NSm might interact with any other components of this vesicle trafficking machinery. Indeed, our mass spectrometry analysis revealed further specific interactions between NSm and two of the NRZ complex subunits, NBAS and ZW10, and we were able to immunologically validate the interaction with ZW10 via western blotting. Furthermore, inhibition of COPI through BFA treatment resulted in loss of NSm protein expression at the Golgi. Although the exact role of the NRZ complex during BUNV infection is unclear, the interactions of NSm with BNIP1 and ZW10 suggest that NSm may exploit host cell COPI vesicle trafficking pathways to promote Golgi-associated VF formation or specific functions within the VF.

Although studies linking BNIP1 or NRZ complex components to other bunyaviruses are lacking, COPI components have been implicated as host cell factors in certain cases. We have previously reported COPI subunits to be necessary for the efficient replication of Hazara virus, a bunyavirus belonging to the *Nairoviridae* family that shares several key characteristics with the peribunyaviruses. COPI was found to play a role in early and late stages of the replication cycle, with knockdown of COPI significantly reducing viral titers (50). Additionally, we have identified COPI as a host cell factor required for efficient virion production and release in the case of lymphocytic choriomeningitis virus (LCMV) within the *Arenaviridae* family, with inhibitor usage similarly negatively impacting viral titers (35). Reports such as these suggest that reliance on the COPI trafficking pathway may be a common feature of bunyavirus infections, although the precise mechanisms involved remain to be fully elucidated.

Our findings indicate that NSm interacts with host cell components involved in COPI vesicle trafficking, including BNIP1 and ZW10, raising the possibility that NSm may play a role in modulating host transport routes during infection. Interestingly, the NSm protein of tomato spotted wilt virus, a member of the *Tospoviridae* family of the bunyaviruses, has been implicated in cell-to-cell movement of viral components during infection, suggesting that NSm may function to manipulate host cell transport pathways across different bunyaviruses (51). In the case of BUNV, the localization of NSm to the Golgi-associated VF may indicate a potential connection between NSm and vesicular transport at the VF. However, the precise mechanistic details and functional significance of the interactions between NSm and these host cell protein interactions remains unknown, and while knockdown of BNIP1 significantly reduced BUNV gene expression and virus production, future studies will be necessary to establish whether this directly reflects a role for BNIP1 at the VF. It would be particularly informative to determine whether the selective disruption of these interactions using NSm mutants might affect vesicle trafficking, VF formation, or virus production. Additionally, given that BNIP1 is thought to influence apoptosis and mitochondrial dynamics beyond its role as a t-SNARE (45–48), it would be valuable to explore whether these functions also contribute to efficient BUNV replication.

In summary, we developed a recombinant BUNV variant harbouring an epitope-tagged NSm and used this virus to identify a novel virus-host cell interaction with SNARE protein BNIP1. An siRNA knockdown of expression revealed BNIP1 as a key host cell factor, required for efficient BUNV gene expression and virus production. These findings highlight BNIP1 as a potential target for host-directed antiviral strategies, offering greater durability and reduced potential for resistance compared to virus-targeted therapeutics (52).

## Materials and Methods

### Mammalian cell culture

A549 (adenocarcinomic human alveolar basal epithelial cells), BHK-21 (baby hamster kidney clone 21 cells), and BSR-T7 (BHK-21 cells constitutively expressing T7 RNA polymerase) cells were cultured in Dulbecco’s Modified Eagle Medium (DMEM, Sigma), supplemented with 10% fetal bovine serum (FBS) and 1% penicillin/streptomycin (P/S). BSR-T7 cells were additionally treated with G418 every other passage to maintain the T7 RNA polymerase gene. All cell lines were maintained at 37 °C with 5% CO_2_ in a humidified incubator.

### Plasmids

cDNA plasmids pT7riboBUNS, pT7riboBUNM, and pT7riboBUNL, corresponding to the BUNV S, M, and L segments required for rBUNV-WT virus rescue have been previously described (53, 54). The rBUNV-NSm-HA plasmid was generated firstly through the determination of the predicted structure of NSm using AlphaFold 3 to identify unstructured regions likely to be amenable to tag insertion and accessible to antibodies. The HA tag, flanked on either side by GSGG linker sequences, was inserted between residues V439 and V440 of the of the M segment precursor (18, 19) by PCR using the Q5 Site-Directed Mutagenesis Kit (New England Biolabs) and primers for HA tag insertion following the manufacturer’s protocol. Generation of the rBUNVΔNSm plasmid has been previously described (18). Briefly, PCR was utilized as before to delete residues E332-V477 of the M segment precursor, removing the whole of NSm except for the NSm signal peptide required for Gc translocation and cleavage sites necessary for GPC processing (18).

### Generation of rBUNV-WT, rBUNVΔNSm, and rBUNV-NSm-HA

Rescue of recombinant BUNV variants by reverse genetics has been previously described (53, 54). Briefly, BSR-T7 cells were seeded into a 6-well plate at 2 x 10^5^ cells per well and incubated overnight at 37 °C. The following day, cells were transfected with 1 μg each of pT7riboBUNS, pT7riboBUNM, and pT7riboBUNL and 300 ng of T7 RNA polymerase, or with the M segment plasmid replaced with the relevant BUNV-M cDNA mutant plasmid in the case of rBUNVΔNSm and rBUNV-NSm-HA rescue, mixed with TransIT-LT1 Transfection Reagent (Mirus, 2.5 µL per 1 μg DNA) in 200 µL of OptiMEM Reduced Serum Media (Gibco) topped up to 2 mL following addition of transfection mixes to cells. Control samples where the pT7riboBUNL was omitted were included for all virus rescues. Cells were incubated for 4 hours at 33 °C, prior to removal of transfection mixes and addition of 2 mL DMEM supplemented with 2.5% FBS and 1% P/S per well (2.5% FBS DMEM). At five days post-transfection, supernatants were used to infect BHK-21 cells seeded in T25 flasks at 3 x 10^5^ cells per flask the previous day, and lysates were collected for western blot analysis. At two days post-infection, supernatants were harvested and centrifuged at 4000 *g* for 20 minutes, with viral stocks aliquoted and stored at −80 °C. Lysates were collected for western blot analysis, and titers of viral supernatants were determined by plaque assays.

### Viral infections

For all viral infections, cell monolayers seeded the previous day were washed twice with phosphate-buffered saline (PBS) before infection with BUNV at the specified MOI in 2.5% FBS DMEM unless otherwise stated. At one hour post-infection, inoculums were removed, cells were washed twice with PBS, and fresh 2.5% DMEM was added for the remainder of the infection. A mock-infected condition was included alongside each infection as a control.

### Plaque assays for determination of viral titers

BHK-21 cells were seeded into a 12-well plate at 2.5 x 10^5^ cells per well and incubated overnight at 37 °C. Subsequently, cells were infected with 10-fold serial dilutions of virus stock made in serum-free DMEM with a one-hour incubation period. Virus dilutions were removed, and 1 mL of 1:1 ratio of DMEM to 1.6% methylcellulose overlay was added per well. Following a three-day incubation of cells at 37 °C, the overlay was removed and cells fixed using 4% paraformaldehyde (PFA) for fifteen minutes. Subsequently, PFA was removed and cells were washed with PBS, after which cell monolayers were stained using 2% crystal violet in 20% ethanol. Plaques were counted to determine viral titers as plaque-forming units/mL (PFU/mL).

### Western blot analysis

Cells were lysed with 1x radioimmunoprecipitation assay (RIPA) buffer (150 mM NaCl, 1% Nonidet P-40 alternative (v/v), 0.1% sodium dodecyl sulfate (SDS, wt/vol), 0.5% sodium deoxycholate (wt/vol), and 50 mM Tris-HCl (pH 7.4)), supplemented with 1x Halt Protease Inhibitor Cocktail (100x, Thermo Fisher Scientific) for 15 minutes on ice. Subsequently, lysates were collected and proteins were resolved on 15% SDS-polyacrylamide gel electrophoresis (SDS-PAGE) gels, before transfer to polyvinylidene difluoride (PVDF) membranes. Transfer was carried out under semi-dry conditions at 15 V for 30 min using the Trans-Blot Turbo Transfer System (Bio-Rad), after which membranes were blocked for one hour in Odyssey blocking buffer (LI-COR), diluted 1:1 with PBS supplemented with 0.1% Tween 20 (v/v, PBS-T). Subsequently, membranes were incubated overnight at 4 °C with the following primary antibodies diluted in 1:1 Odyssey blocking buffer with PBS-T: anti-BUNV NP (in-house, 1:5000), anti-GAPDH (glyceraldehyde-3-phosphate dehydrogenase, Santa Cruz Biotechnology, 1:7500); anti-HA tag (Thermo Fisher Scientific, 1:500), anti-BNIP1 (Proteintech, 1:500), and anti-ZW10 (Proteintech, 1:500). Following four five-minute washes with PBS-T, membranes were incubated with corresponding secondary antibodies, IRDye 800CW Donkey anti-Goat, IRDye 680CW Donkey anti-Mouse, and IRDye 800CW Donkey anti-Rabbit (LI-COR, 1:10000 in 1:1 Odyssey blocking buffer with PBS-T), before visualization on the LICORbio^TM^ Odyssey M Imaging System. Densitometry analysis was performed using Fiji (55).

### Immunofluorescence and confocal microscopy

A549 cells were seeded onto glass coverslips in a 12-well plate at 1 x 10^5^ cells per well, unless otherwise stated. The following day, cells were infected with BUNV at the specified MOI, and fixed for ten minutes with 4% PFA at the designated timepoint post-infection. Subsequently, cells were permeabilized for ten minutes in PBS supplemented with 0.3% Triton X-100 (v/v), before undergoing three PBS washes, followed by a one-hour incubation in blocking buffer (1% (wt/v) bovine serum albumin (BSA) in PBS). Next, cells were incubated for two hours at RT or overnight at 4 °C with the following primary antibodies diluted in 1% BSA: anti-BUNV NP (in-house, 1:5000), anti-HA tag (Thermo Fisher Scientific, 1:500), anti-BNIP1 (Proteintech, 1:500), anti-GM130 (Cell Signaling Technology, 1:3200), anti-TOM20 (Proteintech, 1:200), and anti-CANX (Proteintech, 1:200). Following three PBS washes, cells were incubated with corresponding AlexaFluor fluorochrome-conjugated secondary antibodies chicken α-mouse 488, donkey α-goat 594, and donkey α-rabbit 647 (Life Technologies, 1:500 in 1% BSA). Cells were washed three times with PBS before being mounted onto microscope slides using ProLong Gold Antifade Mountant with DNA Stain DAPI (4’6-diamidino-2-phenylindole, Invitrogen). Stained cells were imaged on an LSM 880 Confocal Microscope (Zeiss) and processed using ZEN software and Fiji (ImageJ), with line scan analyses carried out using Fiji (55).

### Co-immunoprecipitation and LC-MS/MS

1.15 x 10^7^ A549 cells were seeded into a T175 flask and incubated overnight at 37 °C. The following day, cells were infected with rBUNV-NSm-HA at an MOI of 0.5. At 24 hpi, cells were lysed for 30 minutes on ice, using 1.5 mL RIPA buffer supplemented with 1.5 µL PI per flask, and lysates were collected. For immunoprecipitation, 200 µL of Invitrogen Dynabeads Protein G in suspension were put in a tube, with beads separated using a magnet. Suspension buffer was removed, and beads were resuspended in 800 µL 0.02% PBS-T and 20 µL HA tag antibody. The tube was rotated for two hours at RT, before subsequent removal of the antibody solution from the beads. Beads were washed three times with 800 µL 0.02% PBS-T, prior to addition of the lysate to the beads. The tube was rotated overnight at 4 °C, before removal of the unbound lysate. Beads were washed four times with 800 µL 0.02% PBS-T, then transferred to a fresh tube. Beads were resuspended in 50 µL 0.02% PBS-T and stored at −80 °C. This process was repeated for mock-infected cells. LC-MS/MS was performed by the University of Bristol Proteomics Facility, where samples were subjected to in-solution proteolytic digestion prior to mass spectrometry analysis.

### Reverse transfection of siRNA

A master mix of 15 µL Lipofectamine RNAiMAX Transfection Reagent (Invitrogen) and 410 µL Opti-MEM was added per well to three wells of a six-well plate, corresponding to conditions of no siRNA, scrambled siRNA, or BNIP1 siRNA. Subsequently, relevant siRNAs (Thermo Fisher Scientific) were added to the negative control and BNIP1 knockdown wells at a final concentration of 50 pmol per well, and transfection mixes were incubated for 20 minutes at RT. Following this, 2 x 10^5^ A549 cells in 10% FBS DMEM were added per well, and cells were incubated with transfection mixes for 24 hours at 37 °C. At 24 hours post-transfection, media containing siRNA and transfection reagent was removed and 2 mL fresh 10% FBS DMEM added to each well. Following a six-hour recovery period, cells were washed twice with PBS prior to infection with WT-BUNV at an MOI of 0.5. At 24 hpi, viral supernatants were harvested and cells were lysed for plaque assays and western blot analysis, respectively.

### BFA treatment assay

A549 cells were seeded into a 6-well plate at 6 x 10^5^ cells per well, and onto glass coverslips in a 12-well plate at 1.5 x 10^5^ cells per well for western blot and immunofluorescence analyses, respectively. Cells were incubated overnight at 37 °C, before subsequent infection with rBUNV-NSm-HA at an MOI of 5. At 1 hpi, cells were treated with 5 µg/mL BFA or DMSO for up to 8 hpi. Subsequently, cells were either lysed for western blot analysis, or fixed for immunofluorescence and confocal microscopy analysis.

### Graphs and statistical analyses

All graphs were plotted using Origin 2024b software (56). Error bars on graphs represent standard deviation. Statistical significance was determined using the Student’s t-test, with *p*-values ≤ 0.05 regarded as significant.

## Supporting information

Supplementary Data Set S1

## Acknowledgements

This work was supported by a University of Leeds PhD studentship awarded to RFW. JF was supported by grant PID2023-149259NB-I00, funded by MICIU/AEI/10.13039/501100011033 and by ERDF/EU.

The authors gratefully acknowledge Dr Ruth Hughes and Dr Sally Boxall of the Bioimaging and Flow Cytometry Facility, Faculty of Biological Sciences, University of Leeds, for their expert assistance and use of the Zeiss LSM 880 Confocal Microscope, funded by Wellcome Trust grant WT104918MA. The authors thank Dr Kate Heesom and colleagues at the University of Bristol Proteomics Facility for performing the LC-MS/MS analyses and providing the raw mass spectrometry datasets used in this study.

## Author contributions

RFW performed all experiments, analyzed the data, and wrote the initial manuscript draft. JNB acquired the funding, analyzed the data, wrote the initial manuscript draft. JF analyzed the data and edited an advanced draft of the manuscript.

## Data availability statement

All data generated or analysed during this study are included in this published article and its Supplementary Information files

