## Supplementary Data for "Interaction of Bunyamwera Virus Non-Structural Protein NSm with Cellular BNIP1 is Required for Efficient Viral Gene Expression and Replication"

Running Title: NSm-BNIP1 interaction in Bunyamwera virus replication

*Corresponding author

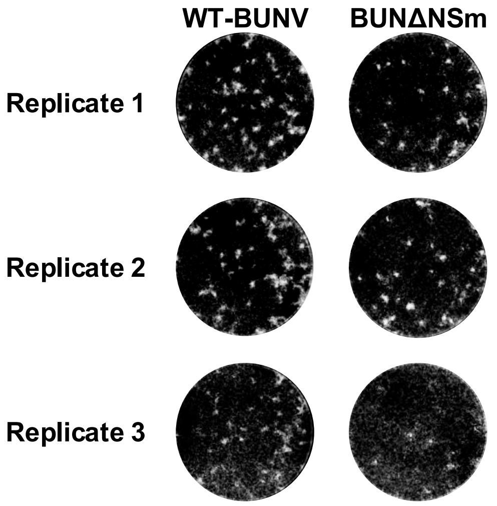


**Supplementary Fig S1.** Representative images of plaque assay wells infected with virus dilutions of $10^{-4}$, corresponding to the graph comparing peak virus yields of rWT-BUNV and rBUNVΔNSm in Fig. 1B.

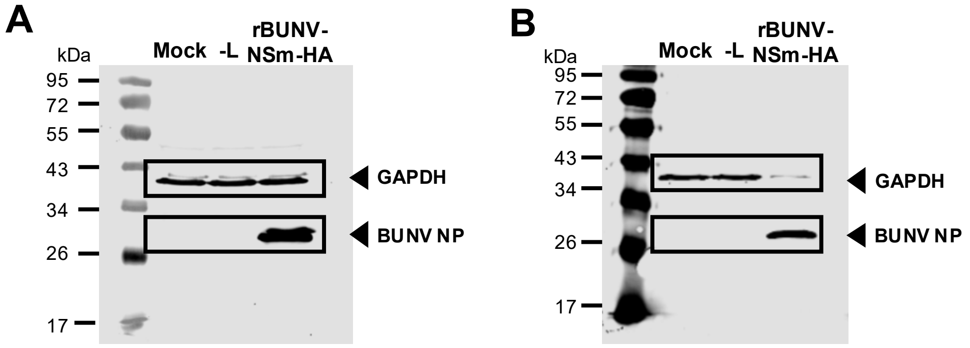


**Supplementary Fig S2.** Uncropped western blots of those displayed in Fig. 2C, with regions shown in the main figure indicated by boxed areas.

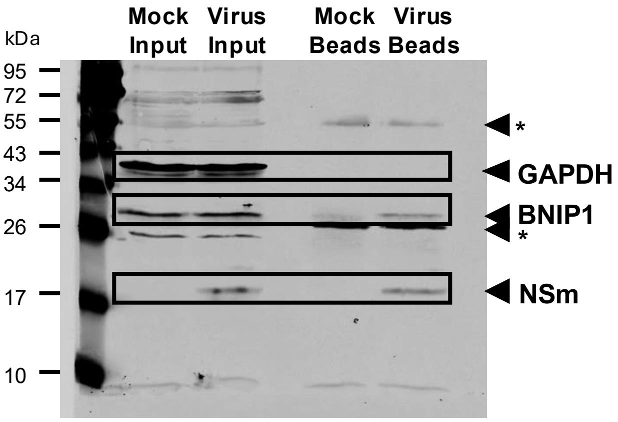


**Supplementary Fig S3.** Uncropped western blot of that displayed in Fig. 4A, with regions shown in the main figure indicated by boxed areas. Asterisks indicate non-specific bands corresponding to antibody heavy and light chains (~50 kDa) and (~25 kDa) from the antibody used for immunoprecipitation.

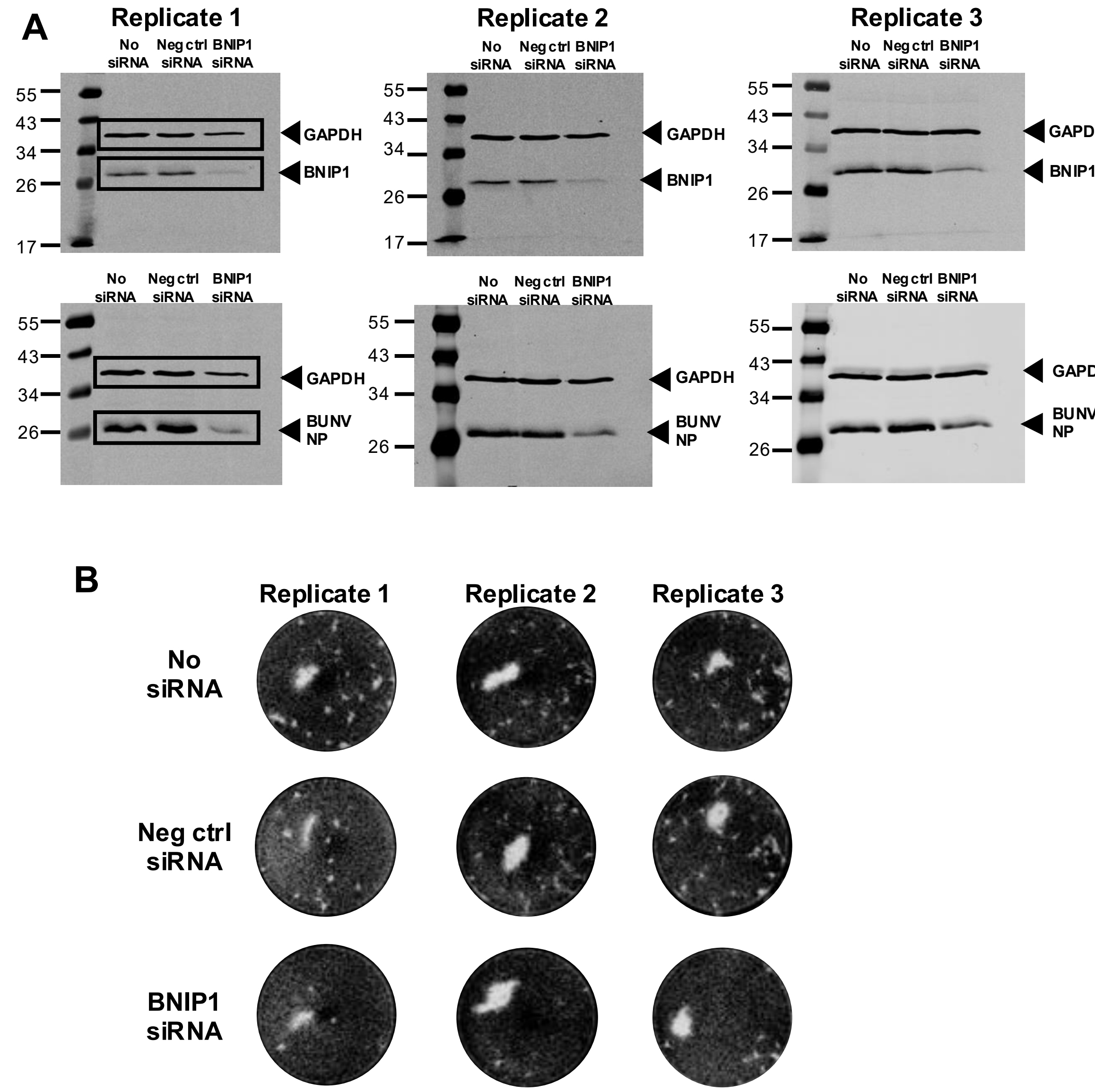


**Supplementary Fig S4.** Effects of siRNA-mediated knockdown of host cell BNIP1 on BUNV gene expression and replication. (**A**) Uncropped western blots of those displayed in Fig. 5A and B, with regions shown in the main figure indicated by boxed areas, alongside uncropped western blots corresponding to the other two biological replicates. (**B**) Representative images of plaque assay wells infected with virus dilutions of $10^{-5}$ for replicate 1, and $10^{-4}$ for replicates 2 and 3, corresponding to the graph comparing viral titers in Fig. 5E.

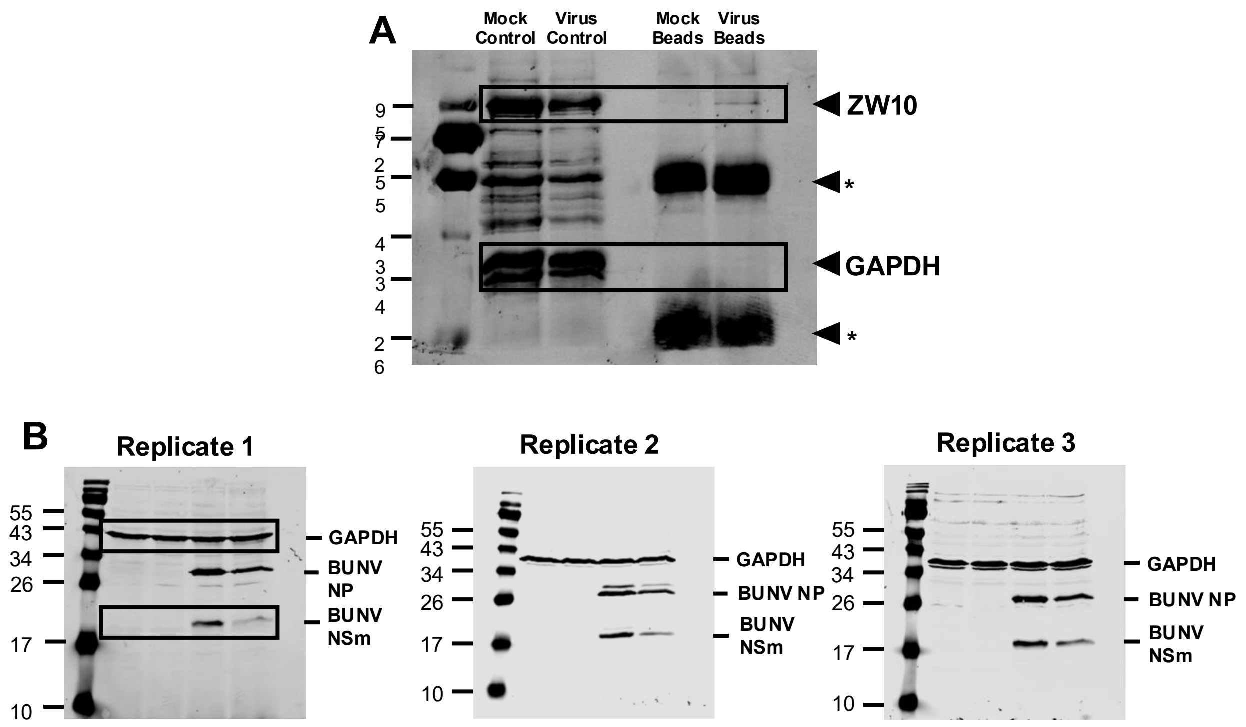


**Supplementary Fig S5.** Analysis of the NSm-ZW10 interaction and the effect of COPI inhibition on NSm protein expression during BUNV infection in mammalian cells. (**A**) Uncropped western blot of that displayed in Fig. 6A, with regions shown in the main figure indicated by boxed areas. Asterisks indicate non-specific bands corresponding to antibody heavy and light chains (~50 kDa) and (~25 kDa) from the antibody used for immunoprecipitation. (**B**) Uncropped western blot of that displayed in Fig. 6C, with regions shown in the main figure indicated by boxed areas, alongside uncropped western blots corresponding to the other two biological replicates.

| Gene | Peak Area Rank | Peak Area | # PSMs | # Unique Peptides |
|---|---|---|---|---|
| ALG1 | 1 | 830014060.875 | 21 | 11 |
| BNIP1 | 2 | 815843529.864583 | 28 | 14 |
| CANX | 3 | 802892685.208333 | 22 | 15 |
| SRSF1 | 4 | 688937643 | 2 | 1 |
| TMEM109 | 5 | 684565750 | 1 | 1 |
| NPC1 | 6 | 325738097.1875 | 1 | 1 |
| SCAMP2 | 7 | 309151510.25 | 3 | 2 |
| POMK | 8 | 251739886.6875 | 8 | 8 |
| TM9SF1 | 9 | 232744882.458333 | 5 | 4 |
| TM9SF2 | 10 | 227749698.875 | 17 | 7 |
| ENSG00000286268 | 11 | 205004681.5 | 2 | 2 |
| SSR3 | 12 | 187973128.5 | 3 | 1 |
| SETDB1 | 13 | 173429791 | 1 | 1 |
| PHB1 | 14 | 170238238.125 | 7 | 6 |
| RNF121 | 15 | 145350561.3125 | 3 | 2 |
| TUBB2B | 16 | 130429300 | 183 | 1 |
| SLC35B2 | 17 | 117355633 | 1 | 1 |
| STT3A | 18 | 113528322.625 | 5 | 3 |
| CLCA2 | 19 | 110619594 | 1 | 1 |
| TNPO1 | 20 | 102767660.791667 | 5 | 5 |

**Supplementary table S1.** Details of the top 20 gene hits by peak area alongside their peptide-spectrum match (PSM) and unique peptide counts from LC-MS/MS analysis of a co-immunoprecipitation, identifying host cell proteins that may interact with NSm in rBUNV-NSm-HA infected A549 cells at 24 hpi at an MOI of 0.5. Before ranking, hits were first filtered by the removal of genes with a peak area of above zero in the non-infected mock samples, as well as those identified as likely laboratory contaminants such as human keratins.
